# Chorioamnionitis as a risk factor for retinopathy of prematurity: an updated systematic review and meta-analysis

**DOI:** 10.1101/291476

**Authors:** Eduardo Villamor-Martinez, Giacomo Cavallaro, Genny Raffaeli, Owais M. M. Mohammed Rahim, Amro M. T. Ghazi, Fabio Mosca, Pieter Degraeuwe, Eduardo Villamor

**Author notes:** Corresponding author (EV). These authors contributed equally to this work (Eduardo Villamor-Martinez, Giacomo Cavallaro).

## Abstract

The role of chorioamnionitis (CA) in the development of retinopathy of prematurity (ROP) is difficult to establish, because CA-exposed and CA-unexposed infants frequently present different baseline characteristics. We performed an updated systematic review and meta-analysis of studies reporting on the association between CA and ROP. We searched PubMed and EMBASE for relevant articles. Studies were included if they examined preterm or very low birth weight (VLBW, <1500g) infants and reported primary data that could be used to measure the association between exposure to CA and the presence of ROP. Of 748 potentially relevant studies, 50 studies met the inclusion criteria (38,986 infants, 9,258 CA cases). Meta-analysis showed a significant positive association between CA and any stage ROP (odds ratio [OR] 1.39, 95% confidence interval [CI] 1.11 to 1.74). CA was also associated with severe (stage ≥3) ROP (OR 1.63, 95% CI 1.41 to 1.89). Exposure to funisitis was associated with a higher risk of ROP than exposure to CA in the absence of funisitis. Additional meta-analyses showed that infants exposed to CA had lower gestational age (GA) and lower birth weight (BW). Meta-regression showed that lower GA and BW in the CA-exposed group was significantly associated with a higher risk of ROP. In conclusion, our study confirms that CA is a risk factor for developing ROP. However, part of the effects of CA on the pathogenesis of ROP may be mediated by the role of CA as an etiological factor for very preterm birth.

## Introduction

Chorioamnionitis (CA) is a major risk factor for preterm birth, especially at earlier gestational age (GA), and a major contributor to prematurity-associated morbidity and mortality [1–5]. The pathogenetic role of CA in the development of adverse outcomes of prematurity, such as bronchopulmonary dysplasia (BPD) [6, 7], necrotizing enterocolitis (NEC) [8], patent ductus arteriosus (PDA) [9, 10], neonatal brain injury [11], or cerebral palsy [12], has been addressed in a number of cohort and case-control studies, which have been summarized in systematic reviews. Nevertheless, it is still controversial whether the effects of CA on neonatal mortality and morbidity are related to infection/inflammation or to the role of CA as an etiological factor for very preterm birth [1–5].

Retinopathy of prematurity (ROP) is a vasoproliferative disorder of the developing retina and a leading cause of childhood blindness around the world [13–19]. Prematurity and postnatal oxygen therapy have consistently been associated with ROP [13–20]. However, ROP is a multifactorial disease, and multiple other modifiable clinical factors have been associated with an increased risk of ROP. These include, among others, hypoxia, hypercapnia, hyperglycemia, exposure to blood transfusions, or poor postnatal weight gain [13–18, 21–26]. In addition, recent experimental and clinical data support the hypothesis that multiple hits of antenatal and postnatal infection/inflammation are involved in ROP etiology and progression [15, 27].

The role of CA as a potential pathogenic factor for ROP has already been the subject of a systematic review and meta-analysis [28]. Mitra et al. [28] included 27 studies (10,590 preterm infants) in their review. They found, in unadjusted analyses, that CA was significantly associated with ROP (any stage, summary risk ratio 1.33, 95% confidence interval [CI] 1.14 to 1.55), and that CA was almost significantly associated with severe ROP (stage ≥3, summary risk ratio 1.27, 95% CI 0.99 to 1.63) [28]. They also carried out subgroup analysis of studies which did not show a significant difference in GA between the CA-exposed and CA-unexposed groups. In this analysis they could not find a significant association between CA and ROP (risk ratio 0.98, 95% CI 0.77 to 1.26). They concluded that CA could not definitively be considered a risk factor for ROP, and that further studies that adjust for potential confounding factors were required [28].

After the publication of the meta-analysis by Mitra et al. [28], more studies assessing the relationship between CA and ROP have been published. Some of these studies are of high methodological quality and included large infant populations. Therefore, in the present study, we aimed to update the meta-analysis of Mitra et al. [28]. We used an extensive search strategy, which included not only studies describing ROP as an outcome after exposure to CA, but also studies that assessed CA as potential risk factor for ROP. In addition, we analyzed the magnitude of the differences in potential confounders, such as GA, birth weight (BW), rate of sepsis, or exposure to antenatal corticosteroids between the infants of the CA and the control group. Finally, we performed a meta-regression in order to investigate the effect of confounders on the association between CA and ROP.

## Methods

The methodology of this study was based on an earlier meta-analysis on the association of CA and PDA [10]. The study was conducted according to the Guidelines for Meta-Analyses and Systematic Reviews of Observational Studies (MOOSE) [29] and the Preferred Reporting Items for Systematic Reviews and Meta-Analysis (PRISMA) [30]. The study is reported according to the PRISMA checklist (S1 File).

### Sources and search strategy

A comprehensive literature search was undertaken using the PubMed/MEDLINE and EMBASE databases from their inception to July 1, 2017. The search terms involved various combinations of the following keywords: “chorioamnionitis”, “intrauterine infection” “intrauterine inflammation”, “antenatal infection” “antenatal inflammation”, “retinopathy of prematurity”, “risk factors”, “outcome”, “cohort”, and “case-control”. No language limit was applied. Additional strategies to identify studies included manual review of reference lists from key articles that fulfilled our eligibility criteria and other systematic reviews on CA, use of “related articles” feature in PubMed, and use of the “cited by” tool in Web of Sciences and Google scholar.

### Study selection

Studies were included if they examined preterm or very low birth weight (VLBW, <1500g) infants and reported primary data that could be used to measure the association between exposure to CA and the presence of ROP. Therefore, we selected studies describing ROP as outcome after exposure to CA, and studies that assessed CA as a potential risk factor for ROP. To identify relevant studies, two reviewers (E.V., G.C.) independently screened the results of the searches and applied inclusion criteria using a structured form. Discrepancies were resolved through discussion or consultation with a third reviewer (P.D.).

### Data extraction

A team of three investigators (G.C., S.G., G.R.) extracted data from relevant studies using a predetermined data extraction form, and a second team of four investigators (E.V.-M., A.G., O.R., P.D.) checked data extraction for accuracy and completeness. Discrepancies were resolved by consulting the primary report. Data extracted from each study included citation information, language of publication, location where research was conducted, time period of the study, study objectives, study design, definitions of CA and ROP, inclusion/exclusion criteria, patient characteristics, and results (including raw numbers, summary statistics and adjusted analyses on CA and ROP where available). Severe ROP was defined as ROP stage ≥ 3.

### Quality assessment

Methodological quality was assessed using the Newcastle-Ottawa Scale for cohort or case-control studies [31]. This scale uses a rating system (range: 0-9 points) that scores three aspects of a study: selection (0-4 points), comparability (0-2 points) and exposure/outcome (0-3 points). Studies were evaluated as though the association between CA and ROP was the primary outcome. Two reviewers (E.V.-M. and E.V.) independently assessed the methodological quality of each study. Discrepancies were resolved through discussion.

### Statistical Analysis

Studies were combined and analyzed using COMPREHENSIVE META-ANALYSIS V3.0 software (Biostat Inc., Englewood, NJ, USA). For dichotomous outcomes, the odds ratio (OR) with 95% confidence interval (CI) was calculated from the data provided in the studies. ORs adjusted for potential confounders were extracted from the studies reporting these data. For continuous outcomes, the mean difference (MD) with 95% CI was calculated. When studies reported continuous variables as median and range or interquartile range, we estimated the mean and standard deviation using the method of Wan et al. [32].

Due to anticipated heterogeneity, summary statistics were calculated with a random-effects model. This model accounts for variability between studies as well as within studies. Subgroup analyses were conducted according to the mixed-effects model [33]. In this model, a random-effects model is used to combine studies within each subgroup, and a fixed-effect model is used to combine subgroups and yield the overall effect. The study-to-study variance (tau-squared) is not assumed to be the same for all subgroups. This value is computed within subgroups and not pooled across subgroups. Statistical heterogeneity was assessed by Cochran’s *Q* statistic and by the *I*^*2*^ statistic, which is derived from *Q* and describes the proportion of total variation that is due to heterogeneity beyond chance [34]. We used the Egger’s regression test and funnel plots to assess publication bias. To explore differences between studies that might be expected to influence the effect size, we performed univariate random-effects meta-regression (method of moments) [35]. The potential sources of variability defined a priori were: CA type (clinical or histological), differences in GA and BW between the infants with and without CA, use of antenatal corticosteroids, mode of delivery, rate of small for gestational age (SGA), rate of premature rupture of membranes (PROM), rate of preeclampsia, rate of early-onset sepsis (EOS), rate of late-onset sepsis (LOS), and mortality. Additional sensitivity analyses were performed excluding studies that included infants with GA >32 weeks. A probability value of less than 0.05 (0.10 for heterogeneity) was considered statistically significant.

## Results

### Description of studies

Of 748 potentially relevant studies, 50 met the inclusion criteria [11, 36–84]. The PRISMA flow diagram of the search process is shown in Fig 1. The included studies evaluated 38,956 infants, and included 9258 CA cases, 3251 cases of all stages ROP, and 2720 cases of severe ROP. The included studies and their characteristics are summarized in S1 Table. None of the studies were designed to primarily examine the association between CA and ROP. In 35 studies, the aim was to examine the outcomes, including ROP, of preterm infants with and without maternal CA. Fifteen studies examined the risk factors for ROP, including maternal CA. Nineteen studies used a clinical definition of CA and 26 studies used a histological definition. In two studies [54, 84], ROP was associated with clinical CA and with histological CA separately. In two studies [52, 62], infants were considered to have CA if they had both clinical and histological CA. In the study of Gray et al. [44] infants were assigned to the CA group if they had clinical or histological CA. Finally, 43 of the 50 studies included infants who were at least <32 weeks GA or had a BW <1500g. One study included infants of <33 weeks [65], five studies included infants up to GA 34 weeks [37, 49, 51, 52, 60], and one study included infants of GA <37 weeks [67].

**Fig 1.**
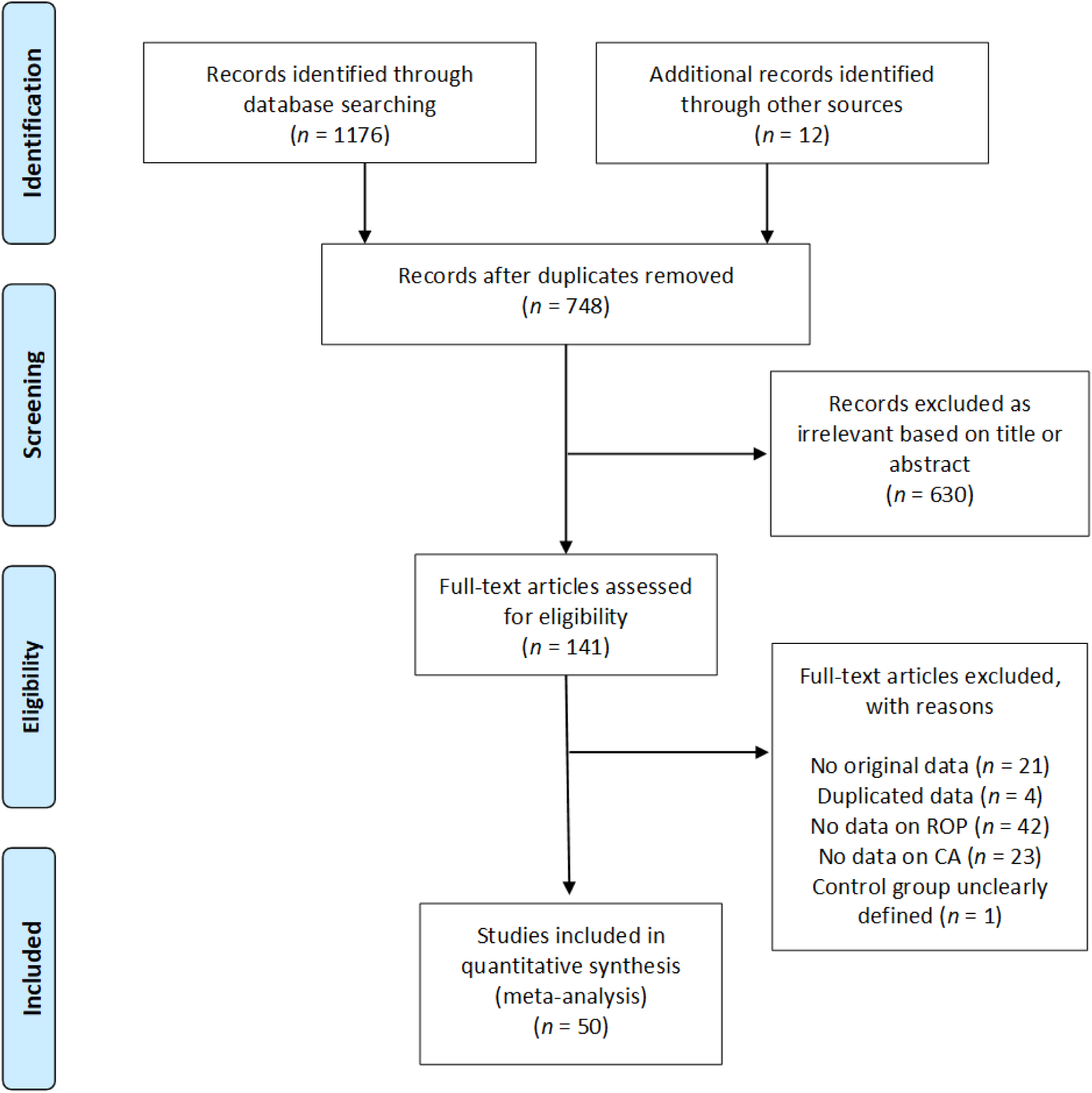
PRISMA flow diagram of search process. CA: Chorioamnionitis; ROP: retinopathy of prematurity

### Quality assessment

The quality of each study according to the Newcastle-Ottawa Scale is summarized in S1 Table. Most (*k* = 40) studies received a quality score of 6 or 7 points. Studies were downgraded in quality most frequently for not adjusting the risk of ROP for confounders (*k* = 42), for not defining ROP clearly (*k* = 24), and for not defining CA clearly (*k* = 16).

### Analysis based on unadjusted data

As shown in Fig 2, meta-analysis showed a significant positive association between all types CA and any stage ROP. The association remained significant for histological, but not for clinical CA. Excluding studies [37, 51, 60, 67] that included older premature infants (GA 32-37 weeks) did not significantly affect the association between CA and any stage ROP (OR 1.34, 95% CI 1.05 to 1.71). Moreover, as shown in Fig 3, meta-analysis showed a significant positive association between all types CA and severe ROP. The association remained significant for both histological and clinical CA. The study of Soraisham, Singhal (65) included older infants (up to 33 weeks GA) and its exclusion did not significantly affect the association between CA and severe ROP (OR 1.62, 95% CI 1.39 to 1.89). Three studies reported on ROP stage ≥1, and meta-analysis demonstrated a significant positive association with CA (S1 Fig). This association became non-significant when a study which included infants up to 34 weeks GA [49] was removed from the analysis (OR 1.89, CI 1.06 to 3.29). Finally, as shown in Fig 4, ROP stage 1-2 was not significantly associated with all types CA, clinical CA, or histological CA. Neither visual inspection of the funnel plot nor the regression test of Egger revealed evidence of publication bias in the analyses of all stages ROP, severe ROP, or stage 1-2 ROP (S2 Fig). There were too few studies (*k* = 3) reporting on stages ≥1 ROP to test for publication bias.

**Fig 2.**
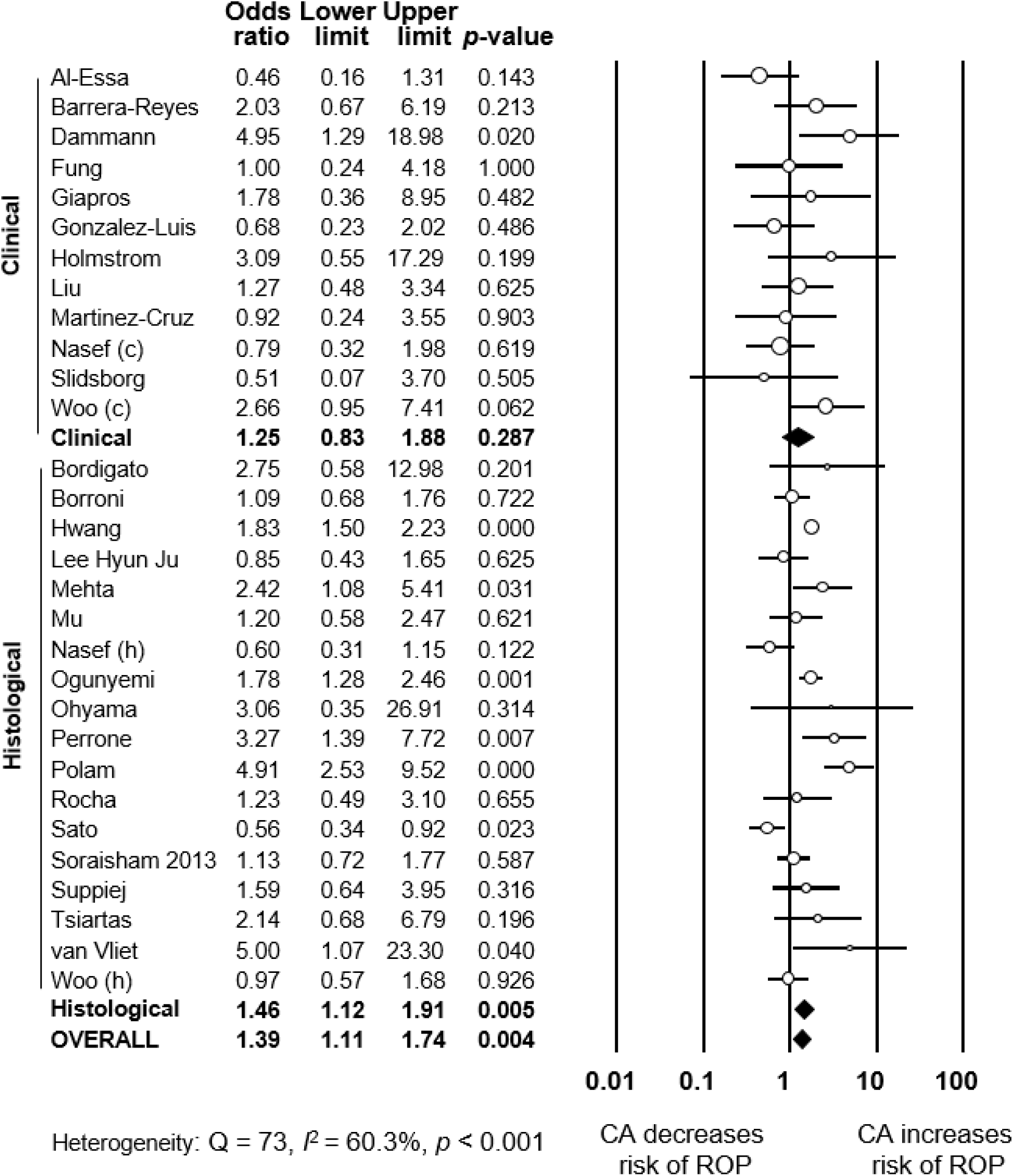
Meta-analysis of CA and risk of any stage ROP. CA: chorioamnionitis; ROP: retinopathy of prematurity; (c): clinical chorioamnionitis; (h): histological chorioamnionitis.

**Fig 3.**
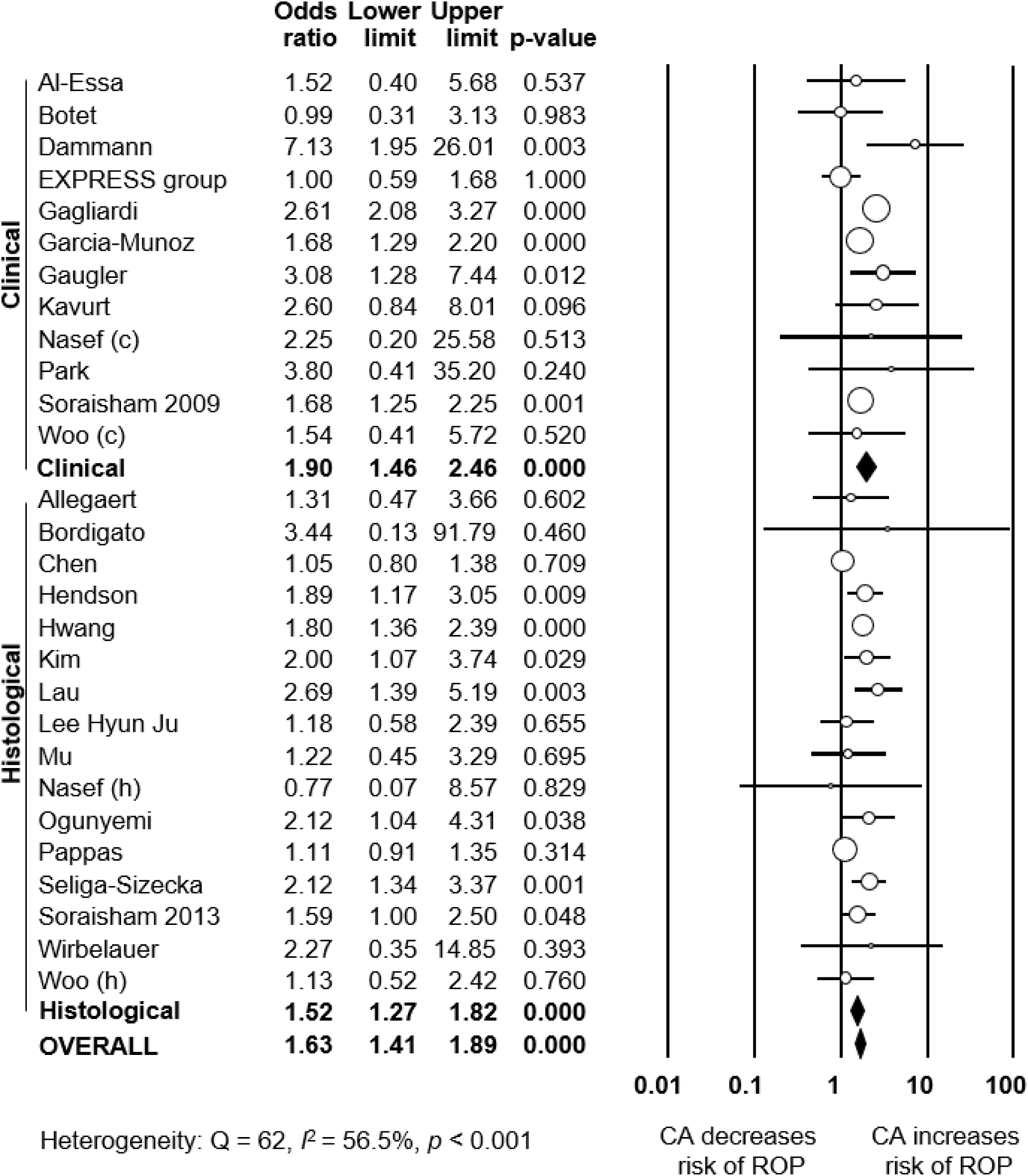
Meta-analysis of CA and risk of severe ROP (stage ≥3) CA: chorioamnionitis; ROP: retinopathy of prematurity; (c): clinical chorioamnionitis; (h): histological chorioamnionitis.

**Fig 4.**
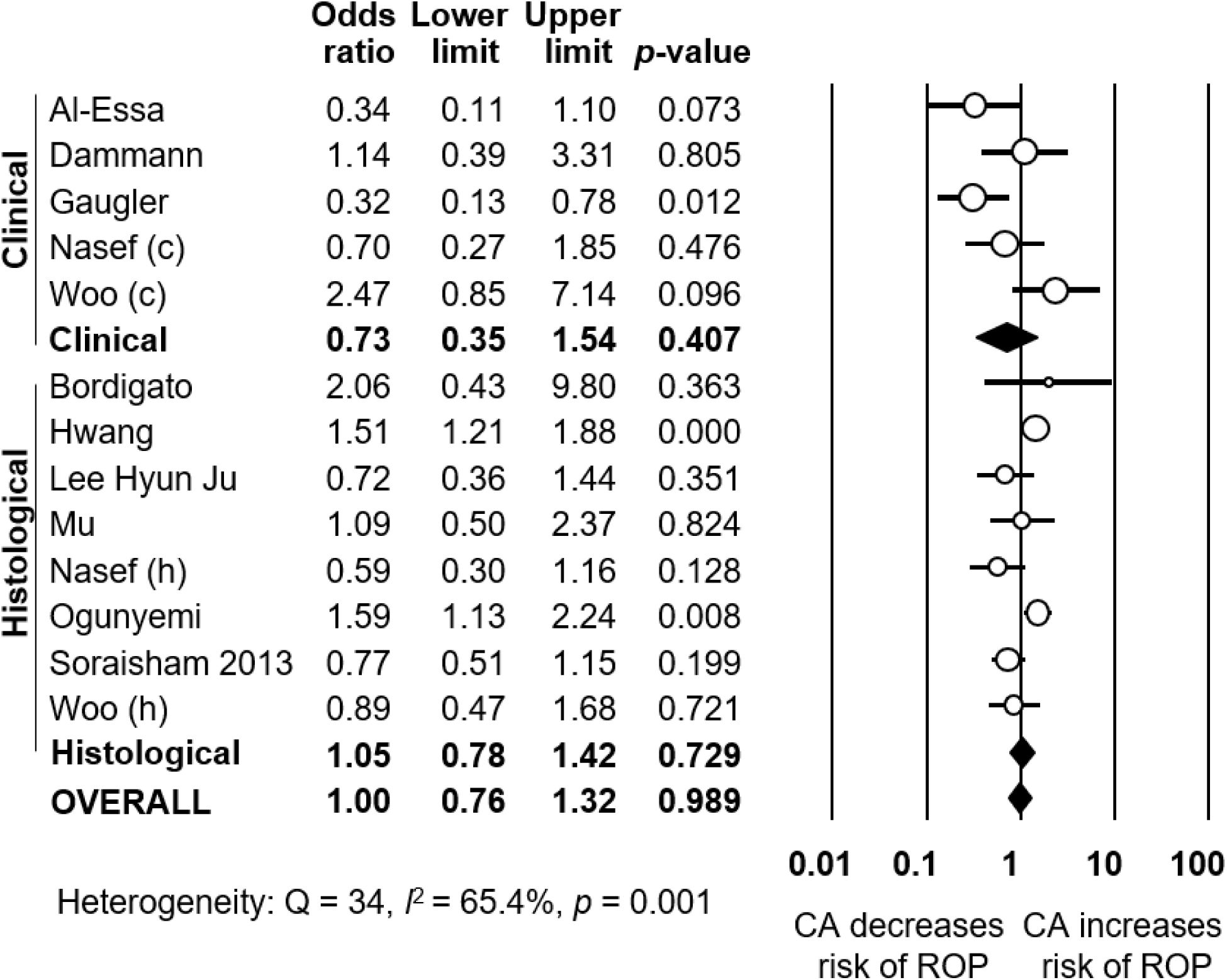
Meta-analysis of CA and risk of stage 1-2 ROP. CA: chorioamnionitis; ROP: retinopathy of prematurity; (c): clinical chorioamnionitis; (h): histological chorioamnionitis. To explore the possible differences in baseline characteristics between the groups exposed and non-exposed to CA, we performed several additional meta-analyses. As summarized in Table 1, infants exposed to CA showed significantly lower GA and BW, significantly lower rates of birth by cesarean delivery, significantly lower rates of SGA, significantly lower rates of preeclampsia, and significantly lower rates of maternal diabetes. Moreover, infants exposed to CA showed significantly higher rates of exposure to antenatal corticosteroids, significantly higher rates of PROM, significantly higher rates of EOS, significantly higher rates of LOS, and significantly higher mortality.

To analyze the possible influence of the GA and BW on the unadjusted association between CA and ROP, we performed meta-regression analyses. These analyses showed that the differences in GA or BW between the CA exposed and non-exposed groups were significantly correlated with the risk of ROP in the CA-exposed group (Table 2). Specifically, we found a significant correlation between an increasing mean difference in GA and a higher CA-associated risk of all stages ROP (Fig 5), stages 1-2 ROP (S3 Fig) and severe ROP (Fig 6). Moreover, we found a significant correlation between an increasing mean difference in BW and a higher CA-associated risk of all stages ROP (S4 Fig), and stages 1-2 ROP (S5 Fig). In contrast, meta-regression could not demonstrate a significant correlation between an increasing MD in BW and a higher CA-associated risk of severe ROP (Table 2).

**Table 1.** Meta-analysis of CA and confounding factors

| Meta-analysis | CA type | *k* | Effect size | 95% CI | Z | *p* | Heterogeneity<br>Q | *p* | *I*<sup>2</sup><br>(%) |
| --- | --- | --- | --- | --- | --- | --- | --- | --- | --- |
| Gestational age (weeks) | Clinical | 7 | MD -0.94 | -1.44 to -0.43 | -3.60 | <0.001 | 158.7 | <0.001 | 96.2 |
|  | Histological | 9 | MD -1.42 | -1.85 to -0.99 | -6.53 | <0.001 | 118.6 | <0.001 | 93.3 |
|  | Any type | 18 | MD -1.15 | -1.44 to -0.85 | -7.61 | <0.001 | 312.5 | <0.001 | 94.6 |
| Birth weight (g) | Clinical | 8 | MD -19 | -72 to 34 | -0.71 | 0.480 | 127.1 | <0.001 | 94.5 |
|  | Histological | 19 | MD -49 | -86 to -11 | 2.57 | 0.010 | 155.5 | <0.001 | 88.4 |
|  | Any type | 29 | MD -34 | -62 to -6 | -2.38 | 0.017 | 294.1 | <0.001 | 90.5 |
| Antenatal Corticosteroids (any dosage) | Clinical | 6 | OR 1.19 | 0.79 to 1.79 | 0.84 | 0.402 | 73.5 | <0.001 | 93.2 |
|  | Histological | 18 | OR 1.28 | 1.00 to 1.64 | 1.96 | 0.050 | 85.3 | <0.001 | 80.1 |
|  | Any type | 25 | OR 1.25 | 1.03 to 1.52 | 2.21 | 0.027 | 162.0 | <0.001 | 85.2 |
| Cesarean section | Clinical | 7 | OR 0.33 | 0.19 to 0.57 | -3.93 | <0.001 | 278.3 | <0.001 | 97.8 |
|  | Histological | 14 | OR 0.34 | 0.23 to 0.49 | -5.52 | <0.001 | 86.8 | <0.001 | 85.0 |
|  | Any type | 22 | OR 0.34 | 0.26 to 0.45 | -7.33 | <0.001 | 368.3 | <0.001 | 94.3 |
| PROM | Clinical | 2 | OR 5.99 | 2.52 to 14.26 | 4.05 | <0.001 | 5.0 | 0.025 | 80.0 |
|  | Histological | 9 | OR 3.37 | 2.21 to 5.16 | 5.61 | <0.001 | 87.6 | <0.001 | 90.9 |
|  | Any type | 11 | OR 3.78 | 2.61 to 5.47 | 7.04 | <0.001 | 99.0 | <0.001 | 89.9 |
| Small for gestational age | Clinical | 2 | OR 0.48 | 0.16 to 1.49 | -1.47 | 0.204 | 0.01 | 0.925 | 0.0 |
|  | Histological | 9 | OR 0.32 | 0.18 to 0.56 | -4.04 | <0.001 | 90.0 | <0.001 | 91.1 |
|  | Any type | 11 | OR 0.35 | 0.22 to 0.55 | -4.44 | <0.001 | 90.1 | <0.001 | 88.9 |
| Preeclampsia | Histological | 5 | OR 0.20 | 0.16 to 0.25 | -13.20 | <0.001 | 1.4 | 0.844 | 0.0 |
| Early onset sepsis | Clinical | 7 | OR 4.51 | 3.29 to 6.19 | 9.36 | <0.001 | 6.1 | 0.409 | 2.1 |
|  | Histological | 10 | OR 3.68 | 2.53 to 5.36 | 6.81 | <0.001 | 21.2 | 0.012 | 57.6 |
|  | Any type | 17 | OR 4.15 | 3.26 to 5.28 | 11.55 | <0.001 | 27.5 | 0.036 | 41.9 |
| Late onset sepsis | Clinical | 4 | OR 1.30 | 0.86 to 1.98 | 1.23 | 0.219 | 12.4 | 0.006 | 75.9 |
|  | Histological | 11 | OR 1.40 | 1.05 to 1.86 | 2.31 | 0.021 | 75.5 | <0.001 | 86.8 |
|  | Any type | 16 | OR 1.35 | 1.09 to 1.67 | 2.78 | 0.005 | 89.1 | <0.001 | 83.2 |
| Maternal age | Any type | 5 | MD -0.31 | -1.61 to 0.99 | -0.47 | 0.639 | 37.2 | <0.001 | 89.3 |
| Maternal diabetes | Any type | 2 | OR 0.71 | 0.53 to 0.95 | -2.29 | 0.022 | 0.1 | 0.775 | 0.0 |
| Mortality | Clinical | 7 | OR 1.66 | 1.26 to 2.18 | 3.65 | <0.001 | 17.1 | 0.009 | 64.9 |
|  | Histological | 13 | OR 1.47 | 1.15 to 1.87 | 3.17 | 0.002 | 37.3 | <0.001 | 67.9 |
|  | Any type | 21 | OR 1.52 | 1.25 to 1.85 | 4.15 | <0.001 | 81.1 | <0.001 | 75.3 |

CA: chorioamnionitis; *k*: number of included studies; MD: difference in means; OR: odds ratio; PROM: premature rupture of membranes; SGA: small for gestational age;

**Table 2.** Meta-regression of difference in gestational age and difference in birth weight and risk of ROP

| ROP stage | Meta-regression | *k* | Coefficient | 95% CI | Z | *p* |
| --- | --- | --- | --- | --- | --- | --- |
| All stages ROP | Difference in mean GA (per week) | 19 | -0.52 | -0.99 to -0.06 | -2.21 | 0.027 |
|  | Difference in mean BW (per 100 g) | 19 | -0.34 | -0.68 to -0.01 | -2.04 | 0.042 |
| Stages 1-2 ROP | Difference in mean GA (per week) | 7 | -0.58 | -0.96 to -0.21 | -3.06 | 0.002 |
|  | Difference in mean BW (per 100 g) | 7 | -0.36 | -0.57 to -0.15 | -3.39 | 0.001 |
| Severe ROP (stage $\geq 3$) | Difference in mean GA (per week) | 17 | -0.49 | -0.66 to -0.31 | -5.30 | <0.001 |
|  | Difference in mean BW (per 100 g) | 16 | -0.03 | -0.22 to 0.16 | -0.29 | 0.774 |

ROP: retinopathy of prematurity; GA: gestational age; BW: birth weight; *k*: number of included studies; CI: confidence interval

**Fig 5.**
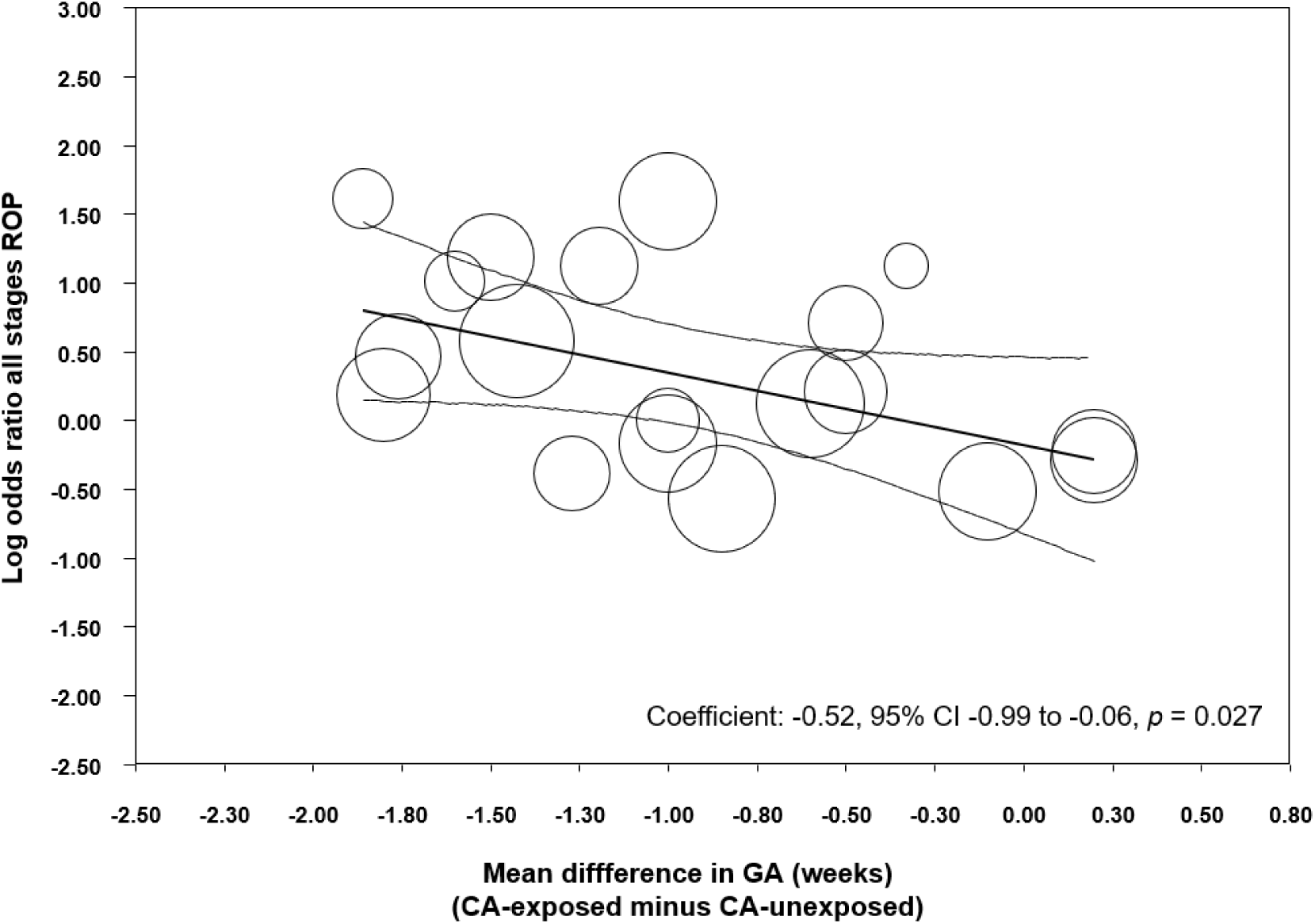
Meta-regression plot of association between CA and all stages ROP controlling for difference in GA between exposed and non-exposed groups. CA: chorioamnionitis; ROP: retinopathy of prematurity; GA: gestational age.

**Fig 6.**
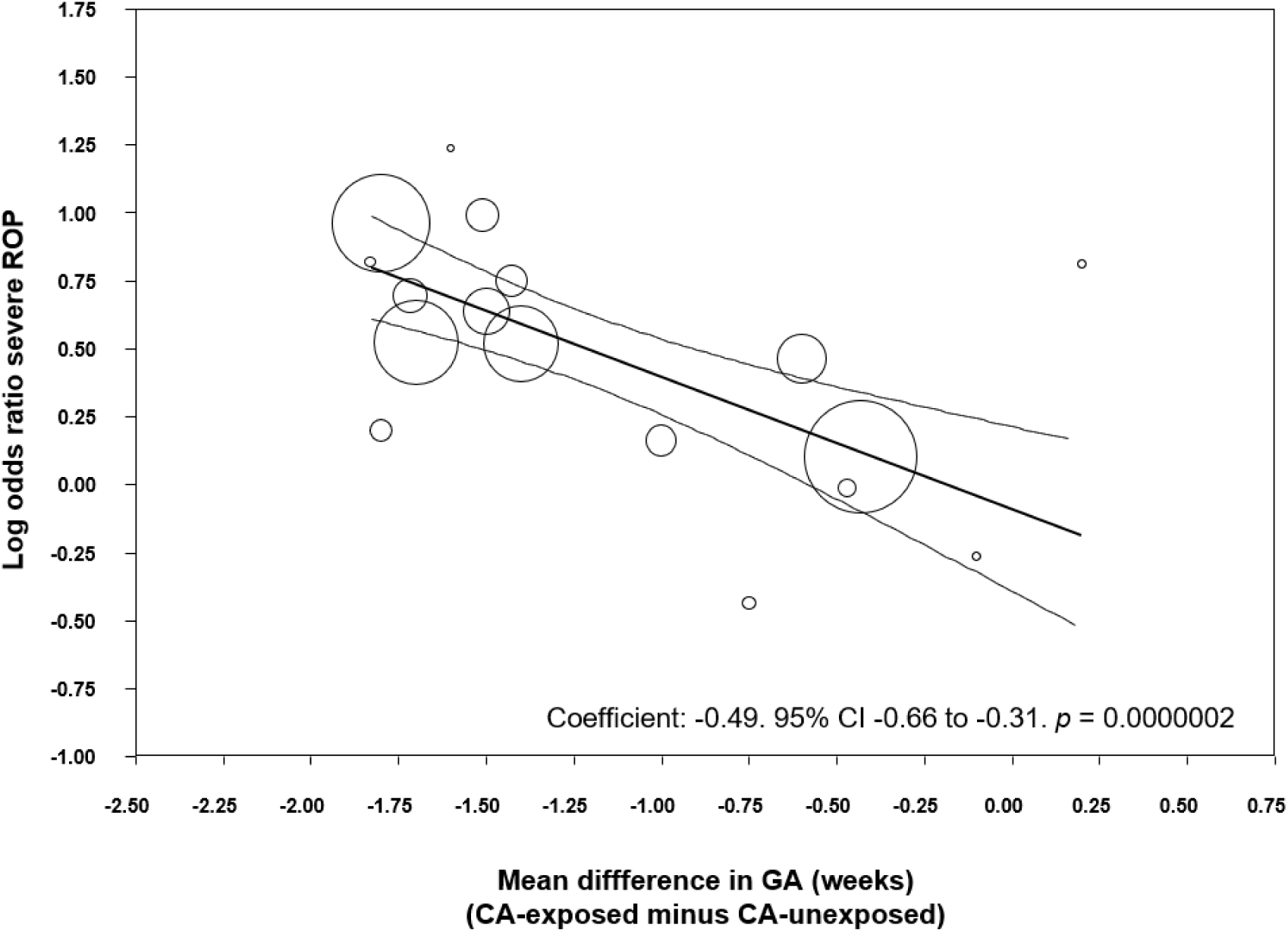
Meta-regression plot of association between chorioamnionitis and severe ROP (stage ≥3) controlling for difference in GA between exposed and non-exposed groups. CA: chorioamnionitis; ROP: retinopathy of prematurity; GA: gestational age.

To eliminate the effect of prematurity as a confounding factor, we carried out a meta-analysis of studies where the mean difference in GA was non-significant (*p* > 0.05). Ten studies met this criterion. As shown in S6 Fig, we could not find a significant association between any type CA and all stages ROP, or any type CA and severe ROP.

In addition, we carried out meta-regression analyses to examine the effect of other covariates on the risk of ROP (S2 Table). We examined the effect of the covariates we predefined on the risk of ROP in the CA-exposed group. We found that an increased risk of EOS in the CA-group significantly correlated with an increased risk of all grades ROP. Moreover, an increased risk of SGA in the CA-group significantly correlated with an increased risk of all grades ROP. Finally, an increased risk of mortality in the CA-group correlated with an increased risk of severe ROP. Other meta-regression analyses of confounders did not show significant associations.

We performed additional analyses aimed at evaluating the role of the presence of fetal inflammatory response (i.e., funisitis) on the development of ROP (S7 Fig). Two studies reported on all grades ROP in infants with histological CA, with or without funisitis. Meta-analysis showed a significant increase in risk of all stages ROP in infants who had funisitis, compared to infants who had CA without funisitis. Two studies reported on severe ROP in infants with funisitis. Meta-analysis showed a significant increase in severe ROP risk in funisitis-positive infants, compared to infants who were only CA-positive (S7 Fig). Finally, when considering any stage ROP and severe ROP together, we observed an increase in risk of ROP in the funisitis-exposed group (S7 Fig). However, additional meta-analysis showed the groups also differed in degree of prematurity. Funisitis-positive infants had significantly lower GA (MD −1.30 weeks, 95% CI −1.37 to −1.23, *p* < 0.0001) than the CA-exposed infants without funisitis.

### Analysis based on adjusted data

Eight studies reported adjusted data on CA-exposure and ROP risk. As described in S3 Table and S4 Table, studies adjusted for different covariates. Meta-analysis of unadjusted data in these studies showed that CA-infants were at a significant risk of all stages ROP (OR 1.73, 95% CI 1.33 to 2.25, S3 Table). Similarly, unadjusted data from these studies showed a significant risk of severe ROP in the CA-group (OR 2.00, 95% CI 1.46 to 2.74, S4 Table). We compared the results of the unadjusted analyses to the adjusted ORs reported in these 8 studies. When using adjusted data, meta-analysis could no longer find a significant association between CA (histological, clinical and any type) and ROP (all stages ROP [S3 Table] and severe ROP [S4 Table]).

## Discussion

Our updated meta-analysis included a greater pool of studies (50 vs. 27) and a larger number of infants (38,956 vs. 10,590) than the meta-analysis of Mitra, Aune (28), but further confirmed their results. We observed a significant positive association between any CA and all stages of ROP. This association was significant for histological but not for clinical CA. In contrast, both clinical and histological CA were associated with severe ROP. Exposure to funisitis was associated with a higher risk of ROP than exposure to CA in the absence of funisitis. Additional meta-analyses showed that infants exposed to CA had significantly lower GA and lower BW than the infants not exposed to CA. Meta-regression showed that these differences in GA and BW were significantly correlated with a higher risk of ROP in the CA-exposed group. Meta-regression also showed that higher rates of EOS, SGA, and mortality in the CA-exposed group correlated significantly with a higher risk of ROP. In summary, our study confirms that CA is a risk factor for developing ROP. However, part of the effects of CA on the pathogenesis of ROP may be mediated by the role of CA as an etiological factor for very preterm birth.

Assessment of CA as a risk factor for adverse outcomes in (very) preterm infants is hampered by the lack of a ‘normal’ control group. Preterm infants carry a mortality/morbidity risk conferred by whatever condition led to their early delivery [85–87]. Two broad pathological conditions have been identified to lead to very preterm birth: (i) infection/inflammation and (ii) placental dysfunction resulting from vascular malfunction [86, 87]. The first group is associated with CA, preterm labor, PROM, placental abruption, and cervical insufficiency. The second group is associated with hypertensive disorders of pregnancy, and fetal indication/intrauterine growth restriction [86]. In addition to distinct pathophysiological pathways, baseline and clinical characteristics are different between the two groups [86, 88]. Accordingly, our analyses showed that the infants exposed to CA were born significantly earlier (~1.15 weeks), were lighter (~35 g), had a higher rate of exposure to antenatal corticosteroids, had a lower rate of cesarean section, were less often SGA, had a higher rate of PROM, had a higher rate of EOS and LOS, and had a higher mortality. Some of these differences may have had a direct or indirect influence on the development of ROP.

We performed meta-regression analyses to evaluate the potential impact of confounders on the risk of ROP. Meta-regression is a statistical technique which examines the relationship between continuous or categorical moderators and the size of effects observed in the studies [35, 89]. Thus, meta-regression allows for the exploration of more complex questions than traditional meta-analysis. The present meta-regression demonstrated that the studies with higher differences in GA and BW between the CA-exposed and CA-unexposed group were also the studies where infants with CA had a greater risk of ROP. Previous meta-regression analyses found a similar correlation between the differences in GA and BW and the CA-associated risk of BPD [7] and PDA [10].

Additionally, we observed that when the few studies that corrected for GA, BW, and other confounding factors were pooled, they did not show a significant increase in the risk of ROP (S3 Table and S4 Table). Previous meta-analyses on the relationship between CA and BPD [7], or cerebral palsy [12], showed that the positive association observed with unadjusted data was significantly reduced, or became non-significant, when adjusted data were pooled. Moreover, in another meta-analysis, the significant positive association between CA and PDA became a significant negative association when only adjusted data were taken into consideration [10]. As mentioned in the introduction, in their meta-analysis on CA and ROP, Mitra, Aune (28) did not pool the studies with adjusted results but performed a subgroup analysis of studies which did not show a significant difference in GA between the CA-exposed and CA-unexposed group. In this subgroup of studies, CA was not significantly associated with ROP. This finding is confirmed in the present meta-analysis, underlining the idea that the effects of CA on ROP development are, at least in part, related to its ability to induce (very) preterm birth.

That the fetal inflammatory response induced by CA might specifically influence the development of the fetal retina is a biologically plausible hypothesis. It has been suggested that multiple hits of antenatal and postnatal infection/inflammation are involved in ROP etiology and progression [15, 27, 90, 91]. Retinal vascular development occurs while the fetus is in the uterus in a relatively hypoxic environment. When an infant is delivered very prematurely, the retinal development must continue in a hyperoxic environment, even in room air, creating a risk for developing ROP. ROP progresses in two phases. The first phase begins with delayed retinal vascular growth and partial regression of existing vessels and is followed by a second phase of hypoxia-induced pathological vessel growth [92]. Recently, a possible pre-phase in the pathogenesis of ROP has been proposed. This pre-phase would begin prior to birth and would be related to fetal inflammation [15, 91]. Proinflammatory cytokines may exert a direct effect on retinal angiogenesis or sensitize the developing retina to the effects of postnatal oxygen, or other stressors. [15, 91]. After birth, the circulatory instability and fluctuation of oxygen saturation following infection/inflammation may affect the retinal perfusion and lead to increased retinal injury. Our meta-analysis shows that CA is not only a risk factor for ROP but also a risk factor for EOS and LOS. Moreover, meta-regression showed a correlation between the effect size of the CA-ROP association and the CA-EOS association. In addition, the meta-analysis of Been et al. [8] demonstrated that CA was a risk factor for NEC, a complication of prematurity in which inflammation plays an important pathogenic role. Altogether, these data suggest that CA-exposed infants are more prone to be exposed to postnatal infection/inflammation and that this propensity makes these infants more vulnerable to ROP.

CA will not always lead to an inflammatory process extending to the fetal component [93]. Funisitis is considered the histologic counterpart of the fetal inflammatory response syndrome [5, 93]. Our analysis showed that the presence of funisitis increased the risk of developing ROP when compared with CA in the absence of funisitis (S8 Fig). These data support the role of the fetal inflammatory response as etiopathogenic factor for ROP. Nevertheless, the number of studies including data on funisitis was rather limited. In addition, infants with funisitis also presented a significantly lower GA when compared with infants with CA without funisitis. Therefore, as in the case of CA, the effects of funisitis on ROP may be related to mechanisms that involve fetal inflammation but also to mechanisms that induce earlier birth.

A further point of interest is that, as assessed by Dammann et al. [74], risk factor patterns for ROP occurrence and progression might differ. A large proportion of very preterm infants will develop low-grade ROP, while in a small proportion it will progress to high-grade disease [74]. Of note, our meta-analysis could not demonstrate that CA was a significant risk factor for ROP stage 1-2 (Fig 4). Recent clinical data suggest that infection/inflammation mechanisms are mainly related to the more advanced stages of ROP, particularly the so-termed aggressive posterior ROP (APROP) [27, 90].

As mentioned above, the two main etiological groups for very preterm birth are infection/inflammation and placental vascular dysfunction [86, 87]. Two recent meta-analyses have studied the risk of ROP of conditions related to the vascular dysfunction group. Neither Chan, Tang (26) nor Zhu, Zhang (94) could demonstrate that maternal or gestational hypertensive disorders affected the risk of developing ROP. However, they did not analyze the differences in basal characteristics between the group of infants exposed and unexposed to maternal/gestational hypertensive disorders. We speculate that the exposed infants probably had a higher GA than the ‘control’ infants and that this difference may have influenced the risk of ROP.

Limitations of the literature and our systematic review and meta-analysis deserve comment. First, the published literature showed great heterogeneity in definition of CA, and in assessment of confounders. Particularly, criteria for the use of the term clinical CA are highly variable, and recent recommendations propose to restrict the term CA to pathologic diagnosis [95]. Second, none of the included studies evaluated the association between CA and ROP as their main objective. Third, adjusted data were available only from 8 of the 50 studies included in the meta-analysis. In addition, we had to rely on the adjusted analyses as presented in the published reports and the variables which they adjusted for, which were not consistent across studies. On the other hand, the main strengths of the present study are the large number of included studies and the use of rigorous methods, including an extensive and comprehensive search; duplicate screening, inclusion, and data extraction to reduce bias; meta-analysis of baseline and secondary characteristics; and the use of meta-regression to control for potential confounders.

ROP is a multifactorial disease that occurs in the youngest and sickest preterm infants [13–19, 96]. Our data show that CA, particularly when accompanied by funisitis, is a risk factor for ROP, but also a risk factor for being a younger and sicker preterm infant. Clinical and experimental evidence suggests that low GA, oxygen stress, as well as ante- and postnatal infection/inflammation are not only independent risk factors for ROP but also interact beyond additive and even multiplicative patterns [15, 21, 97]. Future preventive and therapeutic strategies aimed to reduce ROP, as well as other complications of prematurity, should be tailored, as much as possible, to the particular pathogenic pathway leading to very preterm birth.

## Conflict of interest and role of funders

The authors declare no conflict of interest, and that they did not receive specific funding for this review.

## Supporting information

**S1 Fig.**
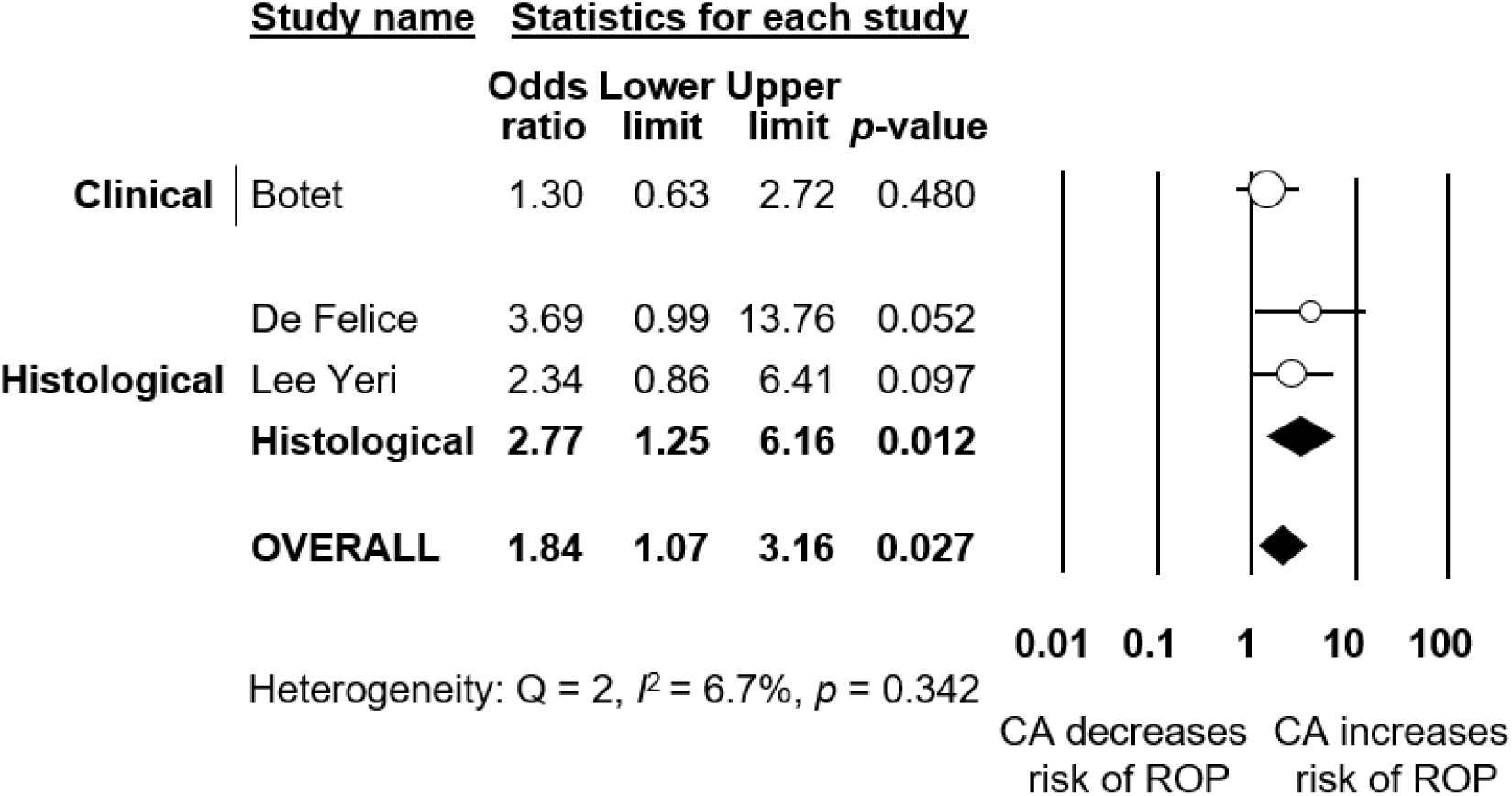
Meta-analysis of CA and risk of stage ≥1 ROP. CA: chorioamnionitis; ROP: retinopathy of prematurity

**S2 Fig.**
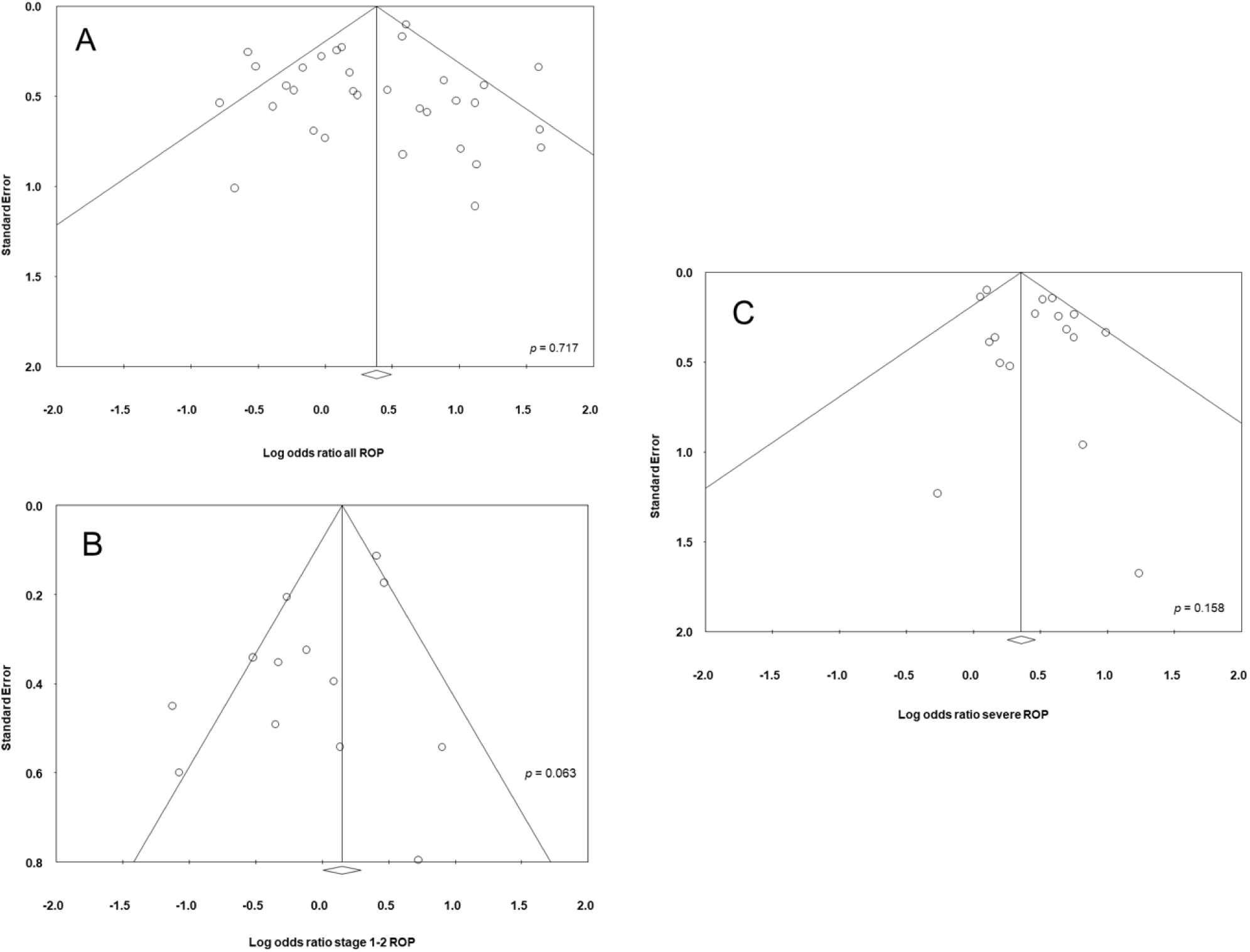
Funnel plot for assessment of publication bias of studies reporting on CA (any type) and all stages ROP (A), severe ROP (B), and stage 1-2 ROP (C). CA: chorioamnionitis; ROP: retinopathy of prematurity

**S3 Fig.**
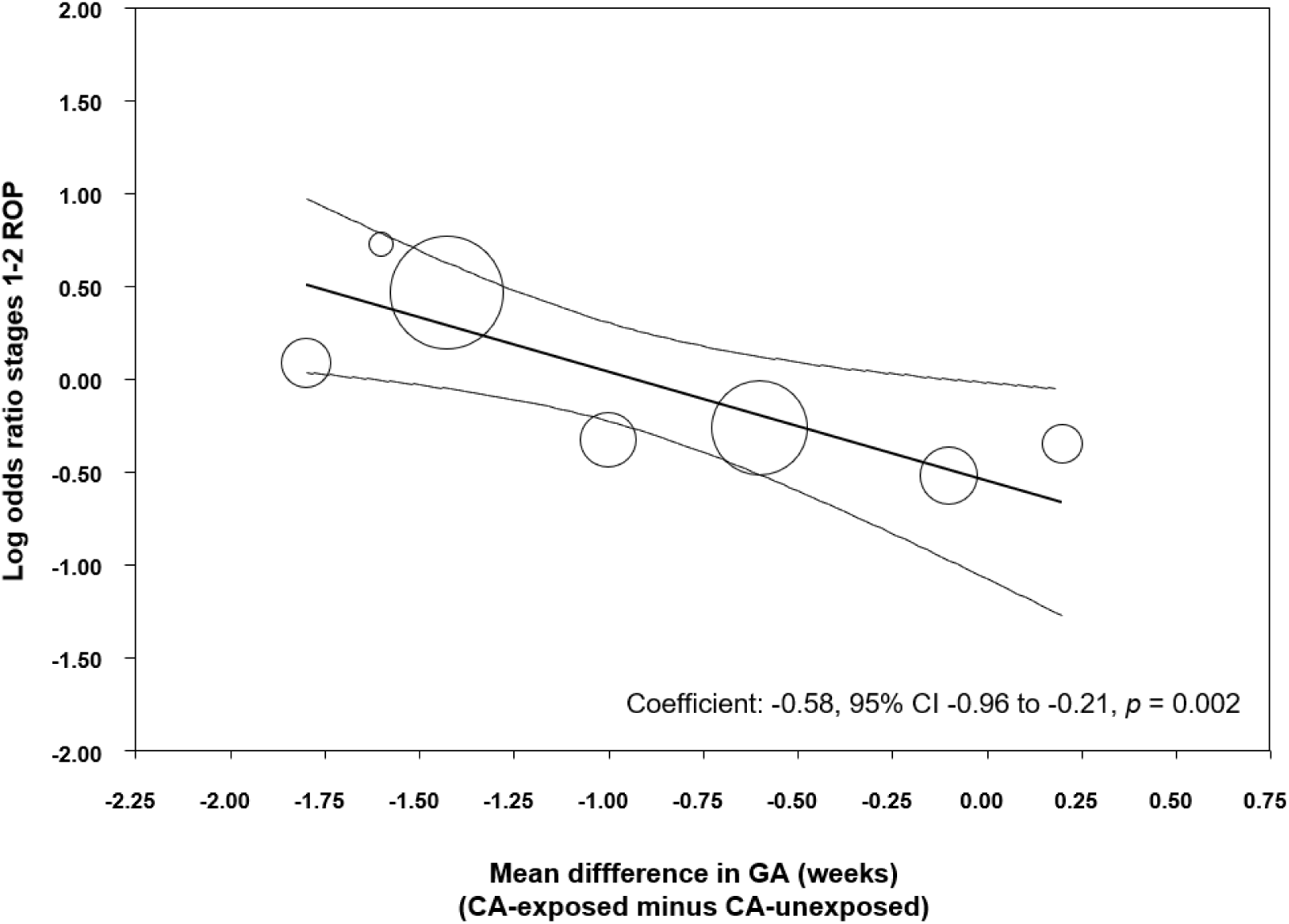
Meta-regression plot of association between CA and stages 1-2 ROP controlling for difference in gestational (GA) between exposed and non-exposed groups. CA: chorioamnionitis; ROP: retinopathy of prematurity; GA: gestational age

**S4 Fig.**
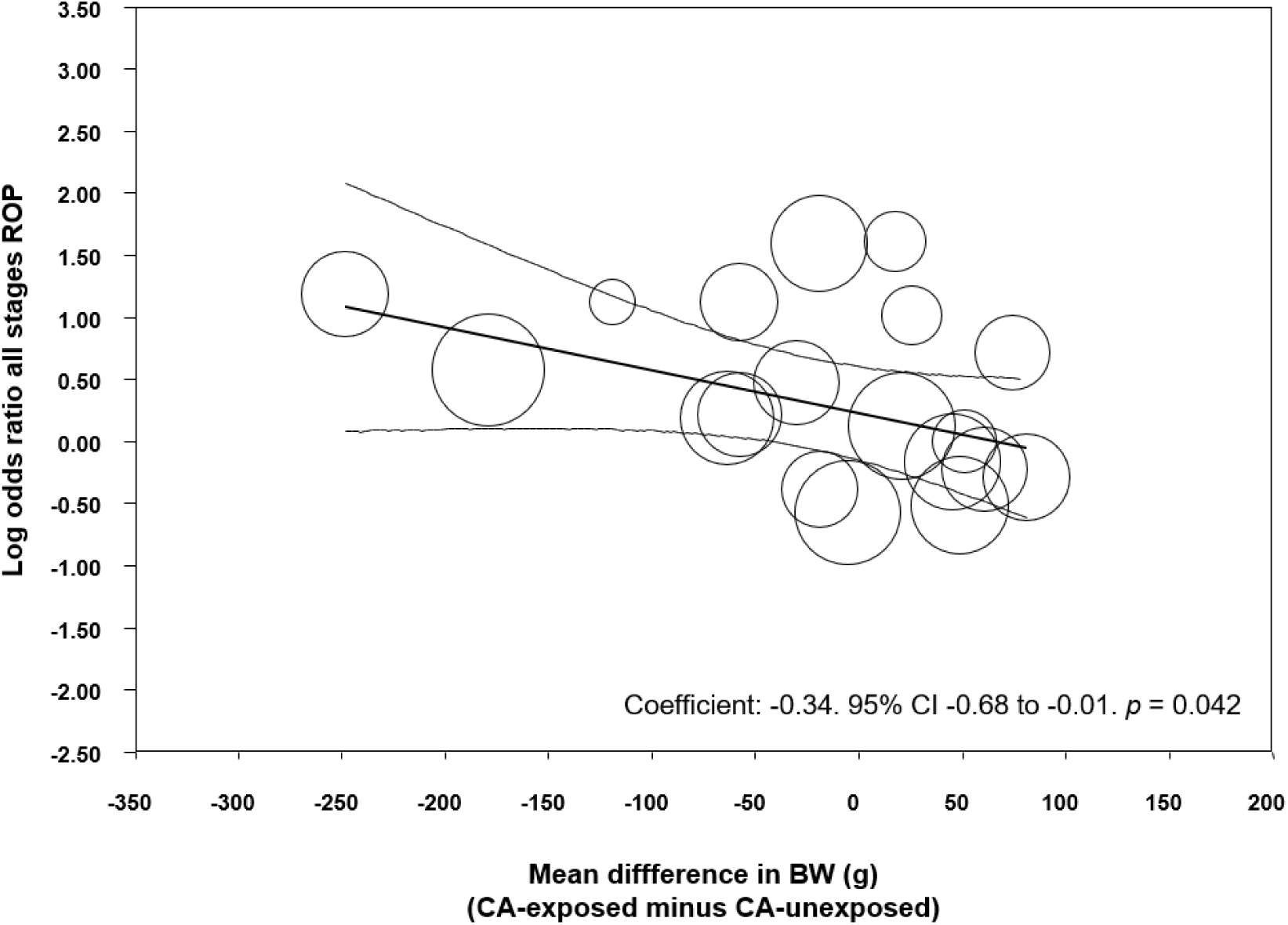
Meta-regression of the relationship between the effect of CA on difference in mean birth weight (BW) and risk of all stages ROP. CA: chorioamnionitis; ROP: retinopathy of prematurity

**S5 Fig.**
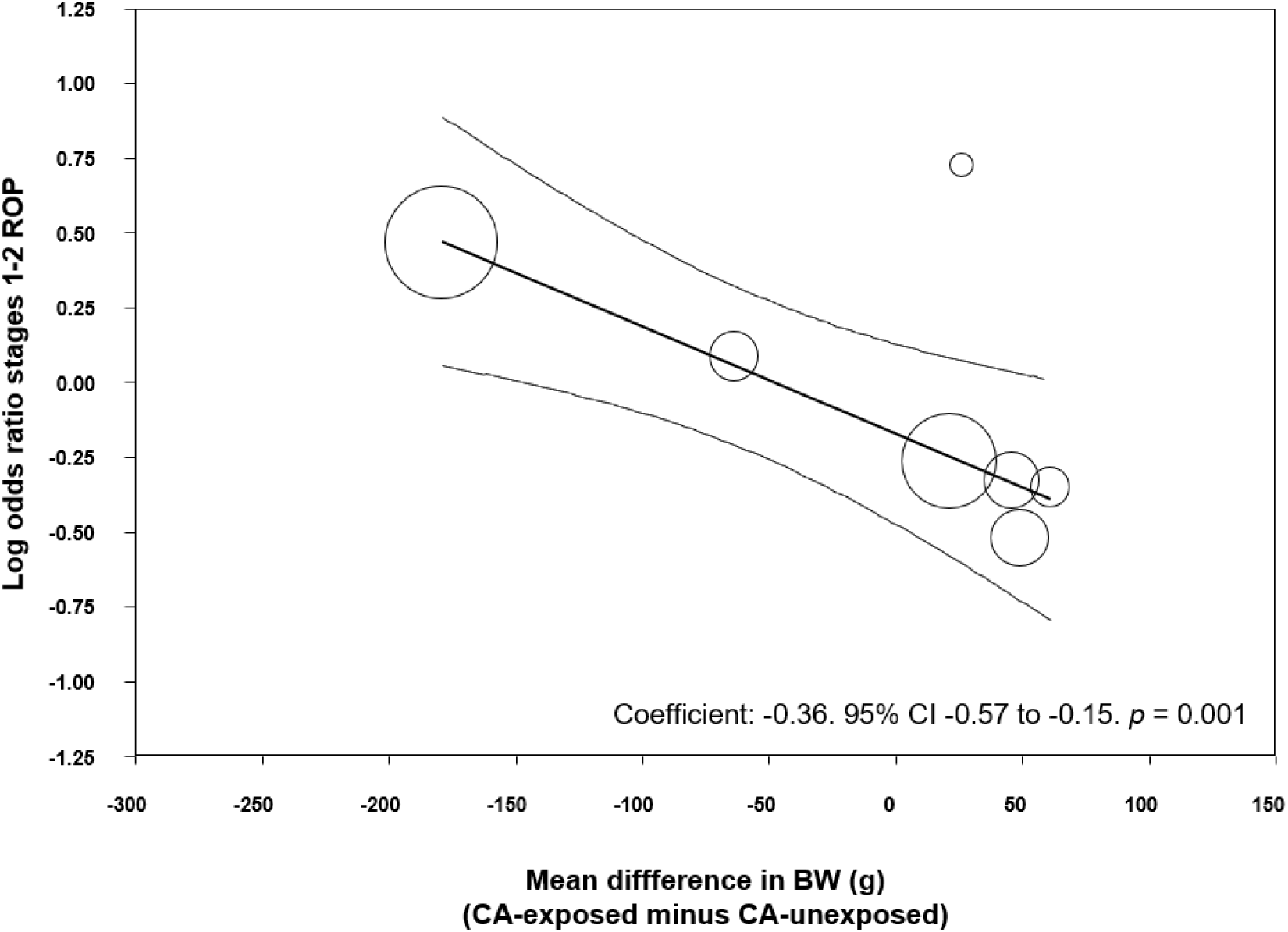
Meta-regression of the relationship between the effect of CA on difference in mean BW and risk of stages 1-2 ROP. CA: chorioamnionitis; BW: birth weight; ROP: retinopathy of prematurity

**S6 Fig.**
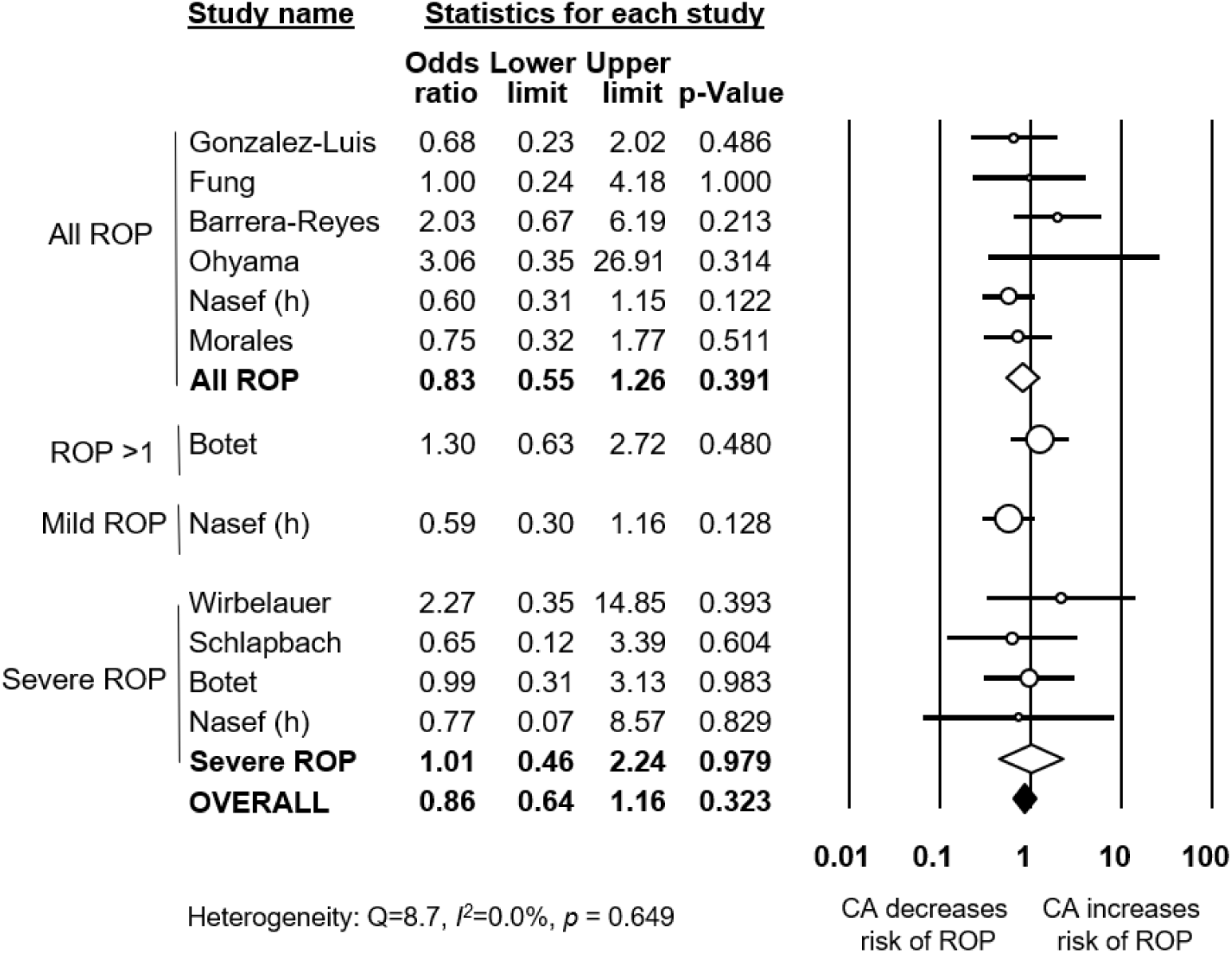
Meta-analysis of CA and risk of ROP, in the subgroup of studies that did not show a significant difference in GA between CA-exposed and CA-unexposed infants. GA: gestational age; CA: chorioamnionitis; ROP: retinopathy of prematurity

**S7 Fig.**
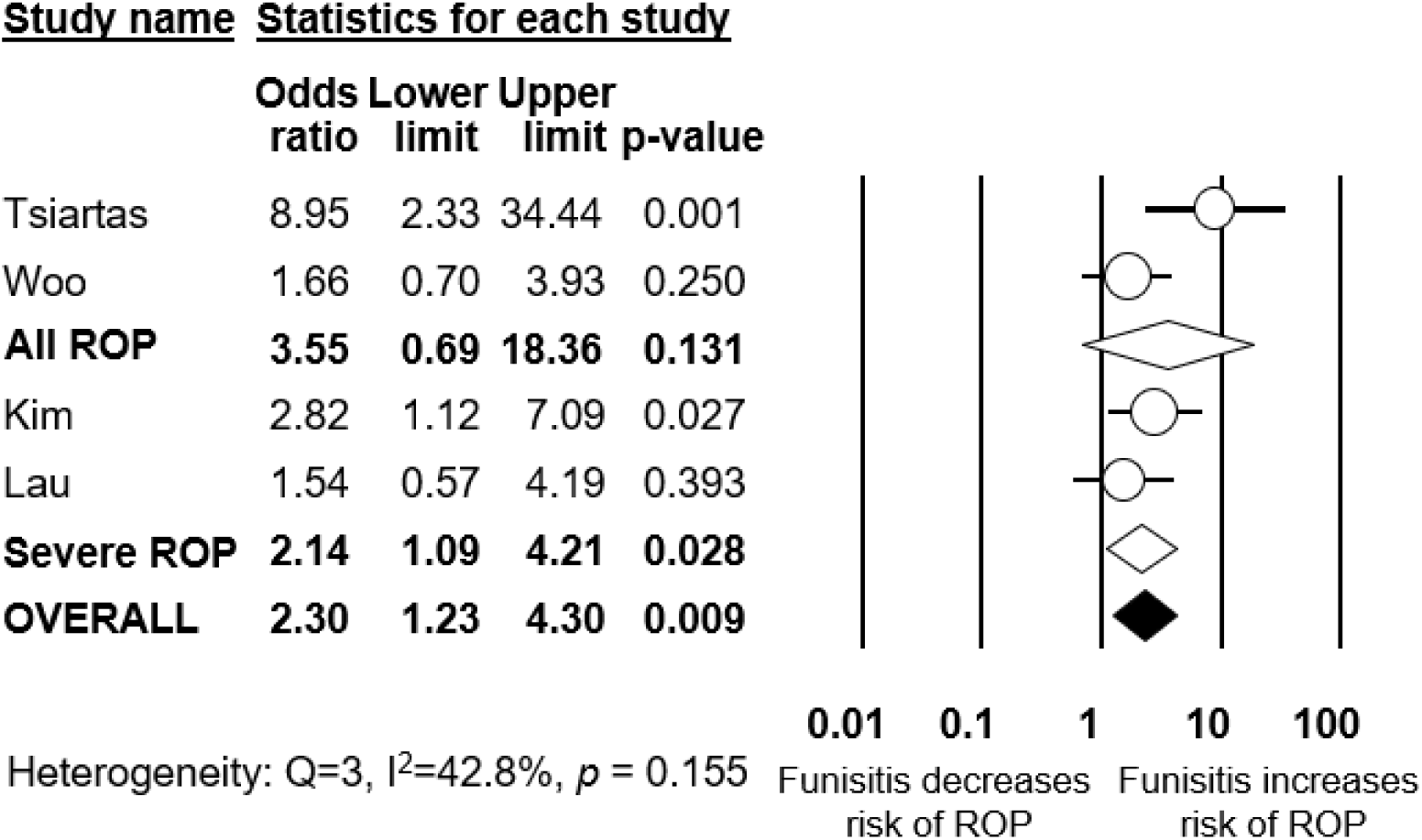
Meta-analysis of funisitis and risk of ROP, compared to CA in the absence of funisitis. CA: chorioamnionitis; ROP: retinopathy of prematurity

**S1 Table.** Synoptic table of all included studies. GA: gestational age; BW: birth weight; ACS: antenatal steroids.

| First author, year | City/region, Country | Design<sup>a</sup> | Perspective | Prosp/Retro<sup>a</sup> | Total infants (centers) | Mean BW (g) | Mean GA (weeks) | Male (%) | ACS (%) | CA category<sup>b</sup> | Definition of CA<sup>c</sup> | Definition of ROP<sup>d</sup> | Newcastle-Ottawa Scale Score<sup>e</sup> |  |  |  |
| --- | --- | --- | --- | --- | --- | --- | --- | --- | --- | --- | --- | --- | --- | --- | --- | --- |
|  |  |  |  |  |  |  |  |  |  |  |  |  | Select | Comp | Outc | Tot |
| Al-Essa, 2000 | Kuwait City, Kuwait | Cohort | ROP | Prosp | 234 (1) | 1145 | 30,2 | 49 |  | CC | NoDes | ICROP | 3 | 0 | 3 | 6 |
| Allegaert, 2003 | Leuven, Belgium | Ca-Co | ROP | Retro | 62 (1) | 827 |  |  | 58 | HC | NoDes | ICROP | 3 | 0 | 2 | 5 |
| Austeng, 2010 (EXPRESS group) | Sweden | Cohort | CA | Prosp | 497 | 800 | 24,3 | 55 | 71 | CC |  | ICROP | 3 | 2 | 3 | 8 |
| Barrera-Reyes, 2011 | Mexico, Mexico | Cohort | CA | Prosp | 104 (1) | 1071 | 30,0 | 52 |  | CC | Ref | NA | 4 | 0 | 2 | 6 |
| Bordigato, 2010 | Padova, Italy | Cohort | CA | Unclear | 29 (1) | 805 | 26,6 | 59 | 76 | HC | Ref | NA | 4 | 0 | 2 | 6 |
| Borroni, 2013 | Italian ROP study group | Cohort | ROP | Prosp | 421 (25) |  |  | 43 | 79 | HC | NoDes | ICROP | 3 | 0 | 2 | 5 |
| Botet, 2011 | Spain | Ca-Co | CA | Prosp | 328 (12) | 1096 | 28,7 | 54 | 84 | CC | Des | NA | 4 | 0 | 2 | 6 |
| Chen, 2011 | ELGAN study group, USA | Cohort | ROP | Prosp | 1062 (14) |  |  | 54 |  | HCF | Ref | ICROP | 4 | 0 | 3 | 7 |
| Dammann, 2009 | Hannover, Germany | Cohort | ROP | Retro | 73 (1) |  | 28,5 | 48 | 81 | CC | NoDes | NA | 3 | 1 | 2 | 6 |
| De Felice, 2005 | Siena and Brindisi, Italy | Cohort | CA | Prosp | 116 (2) | 977 | 28,1 | 48 |  | HC | Ref | NA | 4 | 0 | 2 | 6 |
| Fung, 2003 | Hong Kong, China | Cohort | CA | Prosp | 72 (1) | 794 | 26,2 | 50 | 83 | CC | Des | NA | 4 | 0 | 2 | 6 |
| Gagliardi, 2014 | Italian Neonatal Network | Cohort | CA | Prosp | 3606 (82) | 938 | 27,4 | 50 | 84 | CC | NoDes | NA | 3 | 2 | 2 | 7 |
|  |  |  |  |  |  |  |  |  |  |  |  |  | Select | Comp | Outc | Tot |
| Garcia-Munoz Rodrigo, 2014 | Spanish Network | Cohort | CA | Prosp | 8330 (53) | 1086 | 28,5 | 52 | 67 | CC | Des | NA | 4 | 2 | 2 | 8 |
| Gaugler, 2002 | Strasbourg, France | Cohort | ROP | Retro | 164 (1) | 1007 |  |  |  | CC | NoDes |  | 4 | 0 | 2 | 6 |
| Giapros, 2011 | Ioannina, Greece | Cohort | ROP | Retro | 189 (1) | 1285 | 29,9 | 53 | 58 | CC | NoDes | ICROP | 3 | 0 | 3 | 6 |
| Gonzalez-Luis, 2002 | Barcelona, Spain | Ca-Co | CA | Retro | 135 | 1131 | 28,5 |  |  | CC |  |  | 4 | 0 | 2 | 6 |
| Gray, 1997 | Brisbane, Australia | Cohort | CA | Unclear | 158 (1) | 954 | 27,0 | 56 |  | HC or CC | Des | ICROP | 4 | 0 | 3 | 7 |
| Hendson, 2011 | Edmonton, Canada | Cohort | CA | Prosp | 484 (1) | 930 | 26,9 | 48 | 83 | HCF | Des | ICROP | 4 | 0 | 3 | 7 |
| Holmstrom, 1996 | Stockholm, Sweden | Cohort | ROP | Retro | 202 (5) |  |  |  |  | CC | NoDes | ICROP | 3 | 0 | 3 | 6 |
| Hwang, 2015 | Korean Neonatal Network | Cohort | ROP | Retro | 2009 (55) | 946 | 28,9 | 50 | 76 | HC | NoDes |  | 3 | 0 | 3 | 6 |
| Kavurt, 2014 | Ankara, Turkey | Cohort | ROP | Prosp | 495 (1) | 1267 | 29,3 | 50 | 55 | CC | NoDes |  | 3 | 0 | 3 | 6 |
| Kim, 2015 | Seoul, Korea | Cohort | CA | Retro | 258 (1) | 1104 | 29,2 | 50 | 81 | HCF | Des | ICROP | 4 | 2 | 3 | 9 |
| Lau, 2005 | Canada | Cohort | CA | Prosp | 1296 (1) | 2068 | 33,2 | 55 | 47 | HCF | Ref | ICROP | 4 | 0 | 3 | 7 |
| Lee Hyun Ju, 2011 | Seoul, Korea | Cohort | CA | Retro | 147 (2) | 785 | 26,5 | 56 | 67 | HC | Ref | NA | 4 | 0 | 2 | 6 |
| Lee Yeri, 2015 | Seoul, Korea | Cohort | CA | Retro | 339 (1) | 1525 | 30,0 | 57 | 84 | HC | Ref | ICROP | 4 | 0 | 3 | 7 |
|  |  |  |  |  |  |  |  |  |  |  |  |  | Select | Comp | Outc | Tot |
| Liu, 2014 | Chongqing medical university | Cohort | ROP | Retro | 1614 (1) |  |  |  |  | CC | NoDes | ICROP | 4 | 0 | 3 | 7 |
| Martinez-Cruz, 2012 | Mexico, Mexico | Cohort | ROP | Prosp | 139 (1) | 779 |  | 40 |  | CC | NoDes | Ref | 3 | 0 | 3 | 6 |
| Mehta, 2006 | New Brunswick, USA | Cohort | ROP | Retro | 164 (1) |  |  |  |  | HC | Ref | NA | 4 | 0 | 2 | 6 |
| Morales, 1987 | Orlando, USA | Ca-Co | CA | Prosp | 86 (1) | 1178 | 29,2 |  |  | HC & CC combined | Des |  | 4 | 0 | 2 | 6 |
| Mu, 2008 | Taipei, Taiwan | Cohort | CA | Prosp | 119 (1) | 1108 | 28,6 | 54 | 45 | HC | Ref |  | 4 | 1 | 2 | 7 |
| Nasef, 2013 | Toronto, Canada | Cohort | CA | Retro | 179 (1) | 938 | 27,1 | 55 | 85 | HC & CC separately defined | Ref | Ref | 4 | 0 | 3 | 7 |
| Ogunyemi, 2009 | Los Angeles, USA | Cohort | CA | Retro | 774 (1) | 1313 | 29,4 |  | 53 | HC | Ref |  | 4 | 0 | 2 | 6 |
| Ohyama, 2002 | Yokohama, Japan | Cohort | CA | Retro | 143 (1) | 1162 | 27,8 |  |  | HC | Ref |  | 4 | 0 | 2 | 6 |
| Pappas, 2014 | USA | Cohort | CA | Prosp | 2390 (16) |  | 24,4 | 51 | 75 | HC | Ref | Ref | 4 | 0 | 3 | 7 |
| Park, 2016 | Seoul, Korea | Cohort | ROP | Retro | 201 (1) | 1767 | 32,1 |  |  | CC | NoDes | ICROP | 3 | 0 | 2 | 5 |
| Perrone, 2012 | Siena, Italy | Cohort | CA | Prosp | 92 (1) | 1007 | 26,8 |  |  | HC | Ref | ICROP | 4 | 0 | 3 | 7 |
| Polam, 2005 | New Brunswick, USA | Cohort | CA | Prosp | 177 (1) | 955 | 26,5 | 53 | 74 | HC | Des |  | 4 | 0 | 2 | 6 |
|  |  |  |  |  |  |  |  |  |  |  |  |  | Select. | Comp. | Outc. | Tot. |
| Rocha, 2006 | Porto, Portugal | Cohort | CA | Retro | 452 (3) | 1499 | 29,4 |  | 65 | HC | Ref | ICROP | 4 | 0 | 3 | 7 |
| Sato, 2011 | Yokohama, Japan | Cohort | CA | Retro | 302 (1) | 1211 | 26,3 | 52 | 61 | HC | Ref | Treat | 4 | 0 | 3 | 7 |
| Schlapbach, 2010 | Zürich, Switzerland | Ca-Co | CA | Retro | 99 (1) | 1244 | 29,1 | 49 | 83 | HC & CC combined | NoDes | NA | 3 | 0 | 2 | 5 |
| Seliga-Siwecka, 2012 | Warsaw, Poland | Cohort | CA | Prosp | 383 (1) | 1338 | 29,2 | 56 | 49 | HC | Ref | Ref | 4 | 0 | 3 | 7 |
| Serenius, 2004 | Sweden | Cohort | ROP | Retro | 140 |  |  |  |  | CC |  | ICROP | 3 | 0 | 3 | 6 |
| Slidsborg, 2016 | Denmark | Cohort | ROP | Retro | 6490 |  |  | 54 |  | CC | NoDes | ICROP | 3 | 0 | 3 | 6 |
| Soraisham, 2009 | Canadian Neonatal Network | Cohort | CA | Prosp | 3094 (24) | 1316 | 28,9 | 53 | 79 | CC | Des | ICROP | 4 | 2 | 3 | 9 |
| Soraisham, 2013 | Regional NICU Southern Alberta, Canada | Cohort | CA | Retro | 384 (1) | 885 | 26,3 | 51 | 86 | HC | Des | NA | 3 | 0 | 2 | 5 |
| Suppiej, 2009 | Padova, Italy | Cohort | CA | Prosp | 104 (1) | 1078 | 28,5 | 46 | 87 | HC | Ref | Ref | 4 | 0 | 3 | 7 |
| Tsiartas, 2013 | Králove, Czech Republic | Cohort | CA | Unclear | 231 (1) | 1975 | 33,0 |  | 56 | HCF | Ref | NA | 4 | 0 | 2 | 6 |
| van Vliet, 2012 | Amsterdam, Netherlands | Cohort | CA | Prosp | 72 (1) | 1117 | 29,0 | 51 | 82 | HC | Des | Ref | 4 | 2 | 3 | 9 |
| Wirbelauer, 2011 | Wuerzburg, Germany | Cohort | CA | Prosp | 71 (1) | 1117 | 0,0 | 52 | 94 | HCF | Ref | Ref | 4 | 0 | 3 | 7 |
|  |  |  |  |  |  |  |  |  |  |  |  |  | Select | Comp | Outc | Tot |
| Woo, 2012 | Seoul, Korea | Cohort | ROP | Retro | 246 (1) | 1257 | 29,2 | 57 | 80 | HC & CC separately defined, HCF | Ref | Ref | 4 | 0 | 3 | 7 |

<sup>a</sup>Abbreviations for study design: Ca-Co: case-control study; Perspective, CA: study analyzed ROP as outcome of chorioamnionitis; Perspective, ROP: study analyzed chorioamnionitis as risk factor for ROP; Prosp: prospective; Retro: retrospective;
<sup>b</sup>Chorioamnionitis category: CC: clinical chorioamnionitis; HC: histological chorioamnionitis; HCF: histological chorioamnionitis with funisitis mentioned separately.
<sup>c</sup>Definition of chorioamnionitis: NoDes: no description; Des: clinical or histological description; Ref: defined according to cited article.
<sup>d</sup>Definition of ROP: ICROP: International Classification of Retinopathy of Prematurity; Ref: defined according to cited article; Treat: laser treatment of ROP; NA: no diagnostic criteria mentioned.
<sup>e</sup>Abbreviations for Newcastle-Ottawa Scale: Select: selection; Comp: comparison; Outc: outcome; Tot: total score.

**S2 Table.** Meta-regression of risk of confounding factors and risk of ROP. Log: logarithm; OR: odds ratio; ROP: retinopathy of prematurity; *k*: number of studies included; CI: confidence interval.

| Meta-regression | ROP stage | *k* | Coefficient | 95% CI | Z | *p* |
| --- | --- | --- | --- | --- | --- | --- |
| **Chorioamnionitis type (clinical/histological)** | All ROP | 30 | 0.15 | -0.35 to 0.66 | 0.58 | 0.560 |
|  | Severe ROP | 28 | -0.21 | -0.52 to 0.09 | -1.36 | 0.174 |
| **Antenatal corticosteroids (log OR)** | All ROP | 13 | 0.36 | -0.07 to 0.80 | 1.64 | 0.102 |
|  | Severe ROP | 17 | -0.08 | -0.63 to 0.46 | -0.30 | 0.761 |
| **Cesarean section (log OR)** | All ROP | 10 | -0.10 | -0.85 to 0.66 | -0.25 | 0.801 |
|  | Severe ROP | 16 | -0.21 | -0.43 to 0.02 | -1.83 | 0.067 |
| **Early onset sepsis (log OR)** | All ROP | 7 | 0.65 | 0.23 to 1.07 | 3.03 | 0.003 |
|  | Severe ROP | 11 | 0.23 | -0.46 to 0.42 | -0.07 | 0.946 |
| **Late onset sepsis (log OR)** | All ROP | 8 | -0.13 | -0.80 to 0.54 | -0.39 | 0.695 |
|  | Severe ROP | 12 | 0.49 | -0.05 to 1.03 | 1.76 | 0.078 |
| **Small for gestational age (log OR)** | All ROP | 5 | 0.58 | 0.16 to 1.00 | 2.70 | 0.007 |
|  | Severe ROP | 9 | 0.41 | 0.03 to 0.79 | 2.10 | 0.036 |
| **Premature rupture of membranes (log OR)** | All ROP | 5 | -0.46 | -1.21 to 0.29 | -1.19 | 0.233 |
|  | Severe ROP | 9 | -0.20 | -0.44 to 0.05 | -1.58 | 0.113 |
| **Mortality (log OR)** | All ROP | 11 | 0.30 | -0.24 to 0.85 | 1.10 | 0.273 |
|  | Severe ROP | 13 | 0.74 | 0.37 to 1.11 | 3.95 | <0.001 |

**S3 Table.**
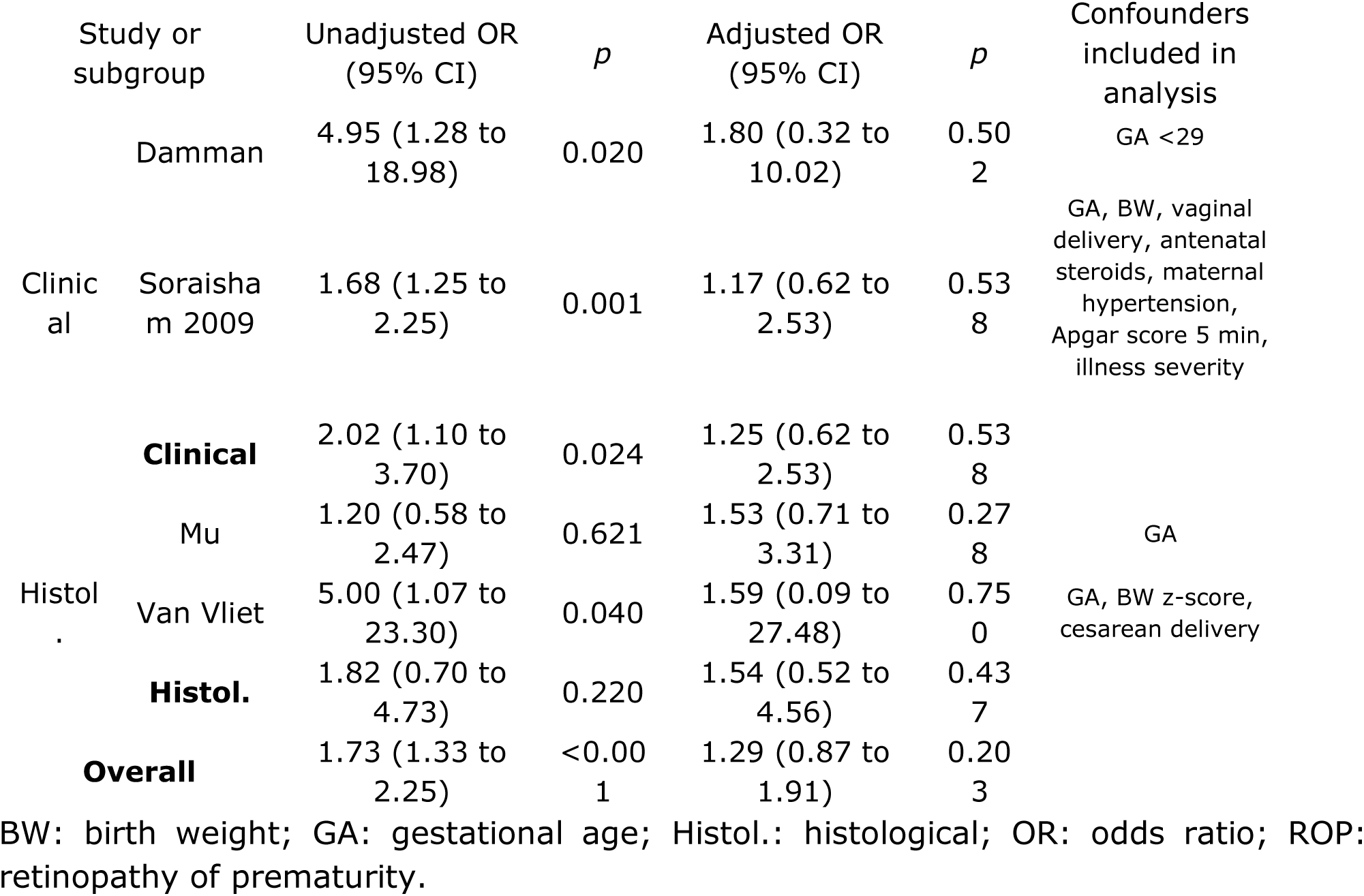
Meta-analysis of crude and adjusted risk of all stages ROP. BW: birth weight; GA: gestational age; Histol.: histological; OR: odds ratio; ROP: retinopathy of prematurity.

**S4 Table.**
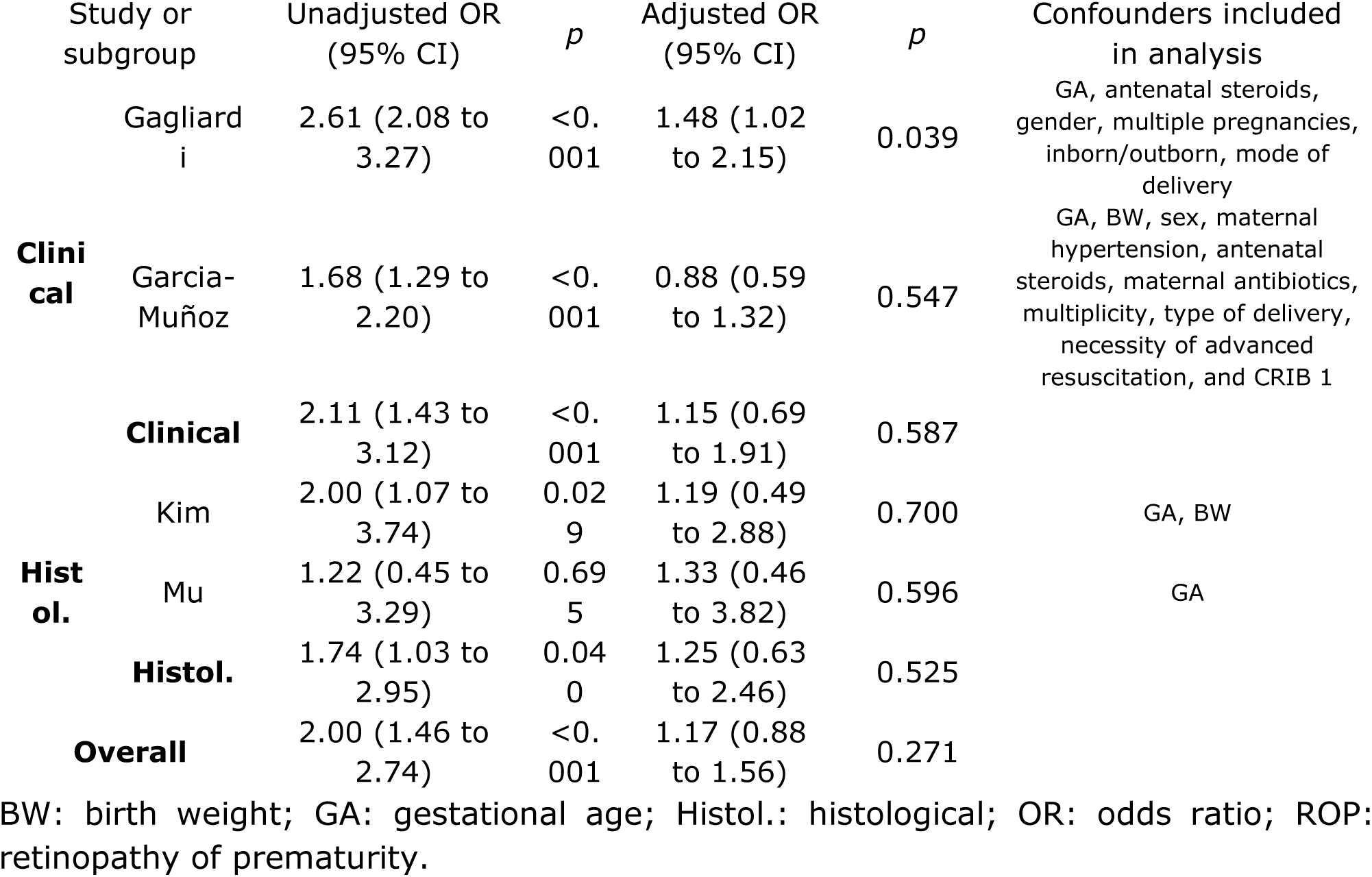
Meta-analysis of crude and adjusted risk of severe ROP (stage ≥3). BW: birth weight; GA: gestational age; Histol.: histological; OR: odds ratio; ROP: retinopathy of prematurity.

S1 File. PRISMA checklist of this review

S2 File. Database of included studies and extracted data

